# Empagliflozin reduces neointimal formation and vascular smooth muscle cell proliferation via the suppression of PDGF-related signaling

**DOI:** 10.1101/2023.08.14.553316

**Authors:** Wei-Jan Chen, Gwo-Jyh Chang, Yu-Juei Hsu, Ying-Hwa Chen

**Author notes:** The first two authors contribute equally to this work. **Address for Correspondence:** Wei-Jan Chen, MD, PhD, Cardiovascular Division, Chang Gung Memorial Hospital, Chang Gung University College of Medicine, Tao-Yuan, Fu-Shin Road no 5, Kwei-Shan, Tao-Yuan 333, Taiwan.

## Abstract

**Objective:** Emerging evidence has documented the beneficial effects of sodium glucose cotransporter 2 (SGLT2) inhibitors on reducing cardiovascular events. Beyond glucose regulation, the mechanisms behind their cardioprotective effects still remained unresolved. This study aims to investigate whether these benefits are mediated by their effects on vascular smooth muscle cell (VSMC) functions.

**Approach and Results:** Treatment of non-diabetic rats with empagliflozin (a SGLT2 inhibitor) attenuated balloon injury-induced neointimal formation in carotid arteries. *In vitro*, treatment of rat VSMCs with empagliflozin reduced platelet-derived growth factor (PDGF)-BB-induced proliferation and migration. Moreover, empagliflozin-treated VSMCs did not undergo apoptosis and cytosis. Notably, treating VSMCs with empagliflozinsuppressed the activation of PDGF-related signaling, such as that related to the phosphorylation of PDGF receptor b (PDGF-Rb), Akt, and STAT3. Furthermore, overactivation of PDGF-related signaling attenuated the inhibitory effects of empagliflozin on VSMC proliferation and migration. The relevant *in vitro* findings were corroborated in the neointima of empagliflozin-treated rats. The fact that minimal SGLT2 was discovered in rat VSMCs and SGLT2 silencing did not alter the effect of empagliflozin supported the SGLT2-independent effect of empagliflozin on VSMC functions.

**Conclusions:** This study highlights the crucial role of PDGF-related signaling in mediating the beneficial effects of empagliflozin on VSMC functions and/or neointimal formation, which are independent of its effects on SGLT2 and glucose metabolism.

---

Emerging evidence has documented the beneficial effects of sodium glucose cotransporter 2 (SGLT2) inhibitors (empagliflozin, dapagliflozin, and canagliflozin) on reducing cardiovascular events in patients with diabetes mellitus.^1–3^ Beyond glucose regulation, the mechanisms behind the cardioprotective effects of SGLT2 inhibitors in these patients still remained unresolved. Potential protective mechanisms, such as lowering blood pressure, improving renal hemodynamics, and alleviating fluid overloading, have been proposed.^4^ Because recent clinical trials have indicated a significant gap between glucose reduction and cardiovascular protection, we hypothesize that these benefits are mediated by the direct cardioprotective effects of SGLT2 inhibitors. Furthermore, we believe that these effects are at least partially independent of the glucose lowering effects of SGLT2 inhibitors.

Neointimal formation is a main pathological feature of atherogenesis, coronary heart diseases, postangioplasty restenosis, and transplantation arteriopathy.^5–7^ Balloon catheter denudation of the endothelium in the carotid artery is commonly used to create a model for studying neointimal formation/restenosis after arterial injury in rats.^8^ In the neointima resulting from balloon injury, vascular smooth muscle cells (VSMCs) exhibit increased proliferation and migration.^5–8^ These proliferative VSMCs could deposit extracellular matrix and accumulate lipids in the intimal layer, contributing to the development of neointimal formation.^5–8^ In this study, we utilized such a model to determine whether SGLT2 inhibitors could inhibit VSMC proliferation and migration and whether such effects are directly caused by SGLT2 inhibitors. The relevant *in vivo* findings were corroborated using VSMCs cultured *in vitro*. The mechanisms and the involved signaling pathways underlying the protective effects of SGLT2 inhibitors against neointimal formation were also investigated.

Growth factors, such as platelet-derived growth factor (PDGF, specially PDGF-BB), play a vital role in regulating the proliferation and migration of VSMCs.^9, 10^ Conceivably, inhibiting PDGF-mediated VSMC proliferation and migration might represent a crucial way for therapeutic intervention in atherogenesis/neointimal formation. PDGF could initiate various biological effects through an activation of downstream signaling.^9, 10^ For instance, PDGF-BB could induce phosphorylation of PDGF receptor b (PDGF-Rb), Akt/glycogen synthase kinase (GSK) 3b, extracellular signal-regulated kinase 1/2 (ERK1/2), and signal transducers and activators of transcription 3 (STAT3).^9, 10^ Accordingly, this study investigated whether PDGF-related signaling might participate in the effects of SGLT2 inhibitors on VSMCs.

## Materials and Methods

### Materials

Most chemicals were purchased from Sigma (St. Louis, MO).

### Rat model of balloon injury

Adult male Wistar rats weighting 350 to 400 g were anesthetized with 3% isoflurane by a ventilator. Angioplasty at right external carotid artery was performed with an inflated 2F Forgarty embolectomy catheter. Empagliflozin (Sigma; 30 mg/kg/day) was administered daily by gavage and balloon injury was performed on day 4. Empagliflozin treatment was continued until rats were euthanized, 14 days after the balloon injury. The uninjured and injured carotid arteries were removed and processed for subsequent biochemical or immunohistochemical analysis. The animal study was approved by the Institutional Animal Care and Use Committee of Chang Gung Memorial Hospital, and performed in accordance with the *Guide for the Care and Use of Laboratory Animals* published by the US National Institutes of Health (the eighth edition, revised 2011).

### Histological and immunohistochemical analyses of rat carotid artery

Carotid artery segments were collected and fixed in 4% paraformaldehyde for 24 h. All the samples were embedded in paraffin, transversely sectioned (5 μm), and stained with hematoxylin-eosin. The neointima/media (I/M) area ratio (an indicator of neointimal formation) was calculated using IMAGE-PRO PLUS (Media Cybernetics, MD). Three discontinuous sections from the middle portion of balloon-injured artery of each rat were measured. Immunohistochemistry was performed by confocal microscopy (Leica-Microsystems, Wetzlar, Germany) using primary antibodies against phospho-(p-Tyr705)/total-STAT3, cyclin D1, and PCNA (Abcam, Cambride, MA) followed by Cy3 (red, Jackson Immuno Research, West Grove, PA)-conjugated secondary antibodies. Nuclei were visualized by 4′,6-diamidino-2-phenylindole (DAPI)-staining. The expression of the target protein was calculated as the protein-occupied area divided by the nuclear area. For each analysis, at least 3 random fields were chosen to observe >5 arterial sections.

### Plasma empagliflozin level

Plasma empagliflozin levels were measured by liquid chromatograph/mass spectrometer (LC/MS-MS) with dapagliflozin (Sigma) as an internal standard. Plasma was placed into a tube containing 500 µg/mL dapagliflozin. The content of each tube was extracted with tert-butyl methyl ether (TBME). Tube contents were mixed and centrifuged for 10 m with 4,000 rcf at 25°C. The supernatant was transferred to another tube, dried under nitrogen gas. The LC-MS/MS analysis was performed using AB SCIEX ExionLC system (Sciex, Canada) coupled with AB SCIEX 5500 QTRAP mass spectrometer (Sciex, Canada). An analytical column (ACQUITY UPLC BEH Phenyl, 1.7 µm, 2.1 x 50 mm; Waters, Milford, MA) was used for the separation. The mobile phase A (water/formic acid, 100/0.1 vol/vol) and mobile phase B (acetonitrile) was degassed and filtered before use. The column flow rate was 0.4 mL/min and the column temperature was maintained at 30°C.

### Cell culture

Rat VSMCs were prepared by enzymatic digestion of the thoracic aortas from adult Wistar rats and cultured in Dulbecco’s modified Eagle’s medium (DMEM) supplemented with 10% fetal bovine serum (FBS) as described elsewhere.^11–14^ Cells used in all experiments were between the fifth and tenth passages. Unless otherwise noted, most experiments were preceded by 48-h serum-deprivation.

### Western blot analysis

Western blot was performed as described elsewhere.^11–14^ Equal amounts of protein sample buffer were sonicated and subjected to sodium dodecyl sulfate-polyacrylamide gel electrophoresis. After being transferred to polyvinylidene fluoride (PVDF) membranes (Stratagene, Netherlands), samples were incubated with primary antibodies against p-Akt(Ser473)/total-Akt, p-ERK1/2(Thr202/Tyr204)/total-ERK1/2, cleaved caspase-3 (Cell Signaling, Beverly, MA), p-STAT3(Tyr705)/total-STAT3, p-PDGF-Rb(Tyr751)/total PDGF-Rb, cyclin D1, PCNA, Bcl-2, and Bax, (Abcam, Waltham, MA), GAPDH, and tubulin (Santa Cruz, Delaware Avenue, CA). Signals were detected by enhanced chemiluminescence (Amersham, Netherlands) and quantified by densitometry. Signal-band intensity was determined relative to that of GAPDH or tubulin.

### Expression vectors and transfection

Wild-type Akt plasmid was cloned by polymerase chain reaction (PCR) using primers: forward (5′-GCGAGATCTATGAACGACGTAGCCA-3′) and reverse (5′-GAAG GATCCTGTGCCACTGGCTGA-3′). The PCR product was subcloned into the pEGFP-N3 vector at the BglIII/BamHI restriction site to generate a GFP-tagged Akt expression vector. Wild-type STAT3 plasmid (GenBank accession no. NM_012747.2) constructed in a pCMV3-C-FLAG vector was purchased from Sino Biological (Chesterbrook, PA). For transient transfection assays, VSMCs at 70-80% confluence were transfected with indicated plasmids using TransIT-LT1 reagent (Life Technologies). The transfection efficiency of this method was approximately 60%.

### Cell proliferation assay

After 48 h of serum deprivation, rat VSMCs were incubated with bromodeoxyuridine (BrdU) from the fifth to the sixteenth h after serum administration. The proliferative activities of VSMCs were quantified by BrdU incorporation using an enzyme-linked immunosorbent assay (ELISA) detecting kit (Roche, Mannheim, Germany) following the manufacturer’s instructions.

### Wound healing assay (Cell migration assay)

After reaching 90% confluence, quiescent VSMCs were treated with 60 ng/mL PDGF-BB and/or 10 μmol/L empagliflozin in serum-free medium for 24 h. A single wound was created by gentle removal of the attached VSMCs with a pipette tip. After 24 h, migrated VSMCs into the wounded area or protruded from the border of the wound were photographed with microscope and counted.

### Real-time quantitative reverse transcription-polymerase chain reaction (RT-PCR)

Total cellular RNA was extracted using TRIzol reagent (Life Technologies, Rockville, MD) and real-time quantitative RT-PCR was performed as described elsewhere.^12, 13^ GAPDH mRNA was used as the internal control.

### Cytotoxicity

Cell cytotoxicity was evaluated by a commercially available CytoScan™ WST-1 Cell Cytotoxicity Assay (ab65473; Abcam).

### Statistical analysis

Data are presented as mean±standard error (SE). Due to the limited number of observations in each group, nonparametric statistical analysis was conducted. Two-group comparisons were performed via unpaired *t*-test, and multiple-group comparisons were performed via the Kruskal-Wallis one-way analysis of variance (ANOVA) followed by Dunn’s method. Interactions among empagliflozin, Akt/STAT3 overexpression, cyclin-D1/PCNA expression, and/or VSMC proliferation/migration were tested with two-way ANOVA. A value of *P* < 0.05 using a two-sided test was considered statistically significant. All statistical analyses were performed using the GraphPad PRISM 5 (GraphPad Software, San Diego, CA).

## Results

### Effect of empagliflozin on neointima formation in balloon-injured rat carotid arteries

This study firstly evaluated the effect of empagliflozin on neointimal formation using balloon-injured model in rat carotid arteries. At 14 days after balloon injury, neointimal formation in the arteries was lower after empagliflozin treatment, as indicated by deceased I/M area ratio, (**Figures 1A and B**). Empagliflozin treatment for 14 days did not have a hypoglycemic effect on any rats (non-fasting blood glucose level: controls=178±12.43; empagliflozin-treated rats=158.56±9.96 mg/dL; standard≈105 mg/dL). The mean plasma empagliflozin concentration after 14 days of treatment was 88.63±23.36 ng/mL (0.157±0.05 μmol/L).

**Figure 1.**
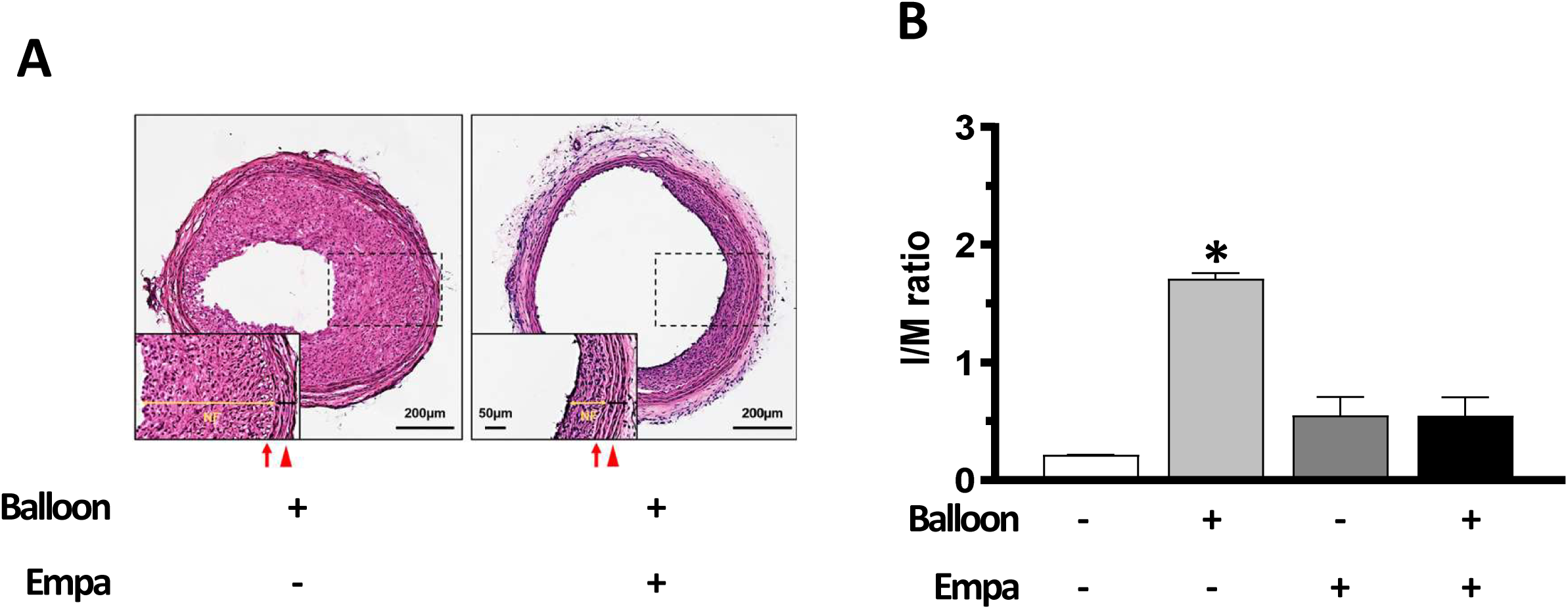
Effect of empagliflozin on neointimal formation in rat carotid arteries after balloon injury. **A.** Representative cross-sections of balloon-injured carotid arteries in control and empagliflozin-treated rats. Arrows denote the internal elastic lamina, and arrowheads denote the external elastic lamina. **B.** Quantitative analysis of neointimal formation by measuring intima/media (I/M) area ratios in uninjured and injured carotid arteries of control and empagliflozin-treated rats. Each value was normalized to the level of control and represented the mean±SE of 10 rats. Using one-way ANOVA, the different symbol (blank, *) represents the significant differences among groups. Empa=empagliflozin; NF=neointimal formation

### Empagliflozin suppressed PDGF-induced VSMC proliferation and migration

Using an *in vitro* culture model, we firstly compared the effects of various cytokines on VSMC proliferation. In agreement with prior reports,^9, 10^ we found that PDGF-BB was the most powerful stimulator of VSMC proliferation, as indicated by the increased levels of cyclin D and PCNA (two important markers of VSMC proliferation)^14^ in western blot analysis (**Figures 2A and B**). Therefore, we utilized PDGF-BB to evaluate the effect of empagliflozin on VSMC functions in the subsequent experiments. Within a concentration of 10 μmol/L, treating VSMCs with empagliflozin concentration-dependently attenuated the promoting effect of PDGF-BB on VSMC proliferation. However, empagliflozin had no effect on VSMC proliferation under basal condition (**Figure 2C**). Consistently, western blot analysis revealed that concurrent administration of empagliflozin with PDGF-BB alleviated PDGF-BB-induced cyclin D and PCNA expressions in a concentration-dependent manner (**Figure 2D**). Furthermore, empagliflozin even at a concentration of 100 μmol/L had no cytotoxic effect on VSMC (**Supplemental Figure 1A**). Although PDGF-BB might induce VSMC apoptosis, empagliflozin treatment could protect VSMCs from PDGF-BB-promoted apoptosis, as indicated by the changes of proapoptotic proteins (Bax and cleaved caspase-3) and anti-apoptotic protein (Bcl-2) (**Supplemental Figure 1B**).

**Figure 2.**
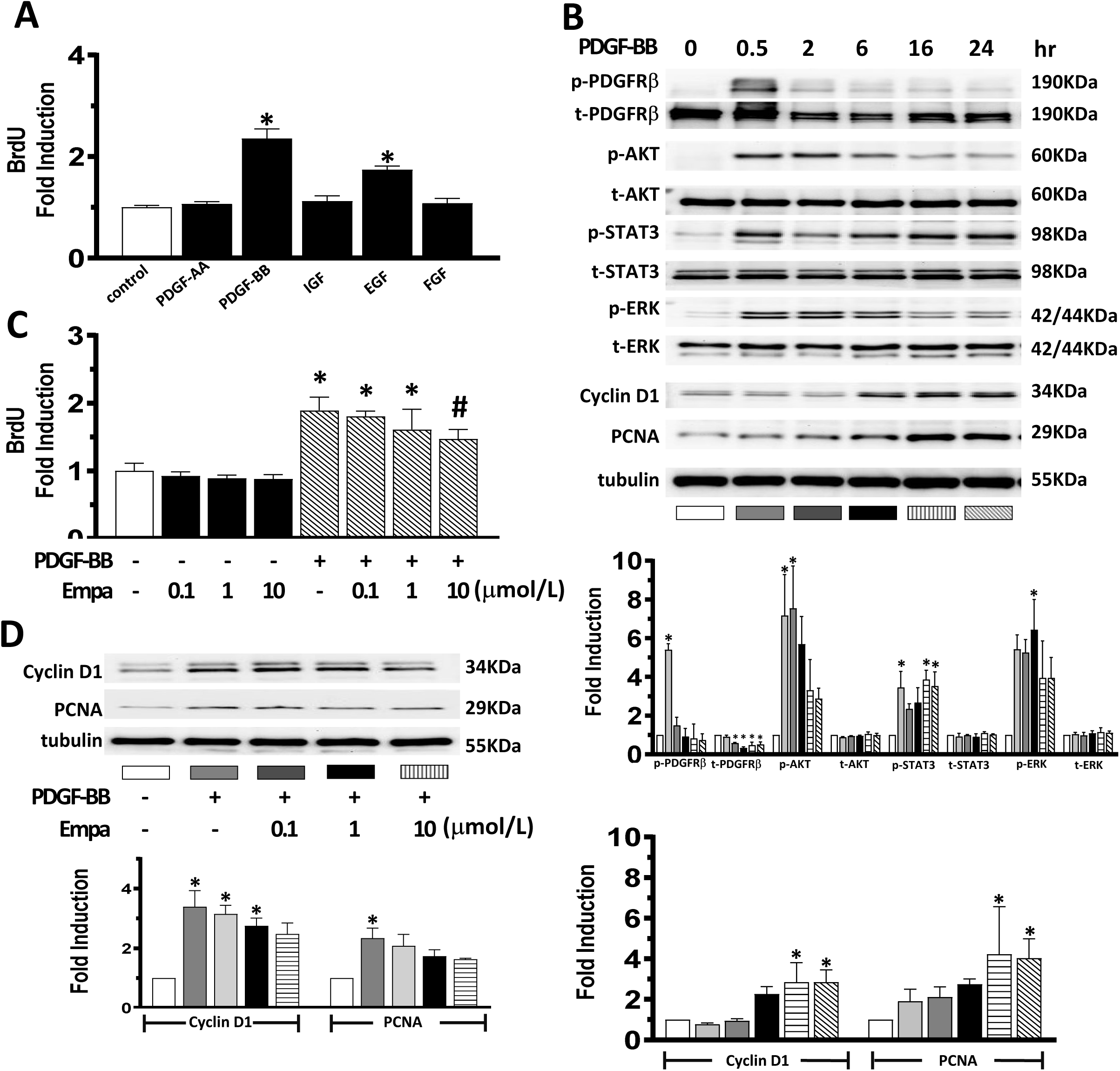
Effect of empagliflozin on PDGF-induced VSMC proliferation *in vitro*. **A.** After 48 h of serum deprivation, VSMCs were treated with indicated cytokines (PDGF-AA: 60 ng/mL; PDGF-BB: 60 ng/mL; IGF: 100 ng/mL; EGF: 50 ng/mL; FGF: 50 ng/mL) for 24 h. The proliferative activity of VSMCs was assayed by BrdU incorporation into VSMCs as described in the method section. The BrdU incorporation level was expressed as the fold change relative to the control condition, which was set at 1.0. IGF=insulin-like growth factor; EGF=epidermal growth factor; FGF=fibroblast growth factor **B.** After 48 h of serum deprivation, VSMCs were treated with 60 ng/mL PDGF-BB for indicated times. The expression of indicated proteins was evaluated by western blot (upper panels). The expression of tubulin was used as an internal control. The relative expression level of each protein was quantified by densitometry and normalized to the control level (lower panels). **C.** After 48 h of serum deprivation, VSMCs were treated with or without 60 ng/mL PDGF-BB and/or indicated concentrations of empagliflozin for 24 h. The proliferative activity of VSMCs was assayed by BrdU incorporation into VSMCs. **D.** After 48 h of serum deprivation, VSMCs were treated with or without 60 ng/mL PDGF-BB and/or indicated concentrations of empagliflozin for 24 h. The expression of indicated proteins was evaluated by western blot (upper panels). The relative expression level of each protein was quantified by densitometry (lower panels). Each value represents mean ± SE of 4 independent experiments. Using one-way ANOVA, the different symbols (blank, *, #) represent the significant differences among groups.

### PDGF-related signaling in empagliflozin-inhibited VSMC functions

We then assessed the involvement of PDGF-related signaling in empagliflozin-induced inhibition of VSMC proliferation. First, time-dependent experiment showed that PDGF-BB promoted the phosphorylation of PDGF-Rb with the maximal effect occurring at 30 min, preceding the changes of p-Akt (30 min to 2 h), p-STAT3 (30 min to 24 h), p-ERK1/2 (6 h), cyclin D1, and PCNA (16-24 h) (**Figure 2B**). As expected, treating VSMCs with empagliflozin concentration-dependently alleviated the effect of PDGF-BB on p-PDGF-Rb, p-Akt, p-STAT3, and p-Erk expression (**Figure 3A**). To further confirm the crucial role of PDGF-related signaling in empagliflozin-induced inhibition of VSMC proliferation, gain-of-function study was applied. We manipulated PDGF-BB-related signaling via transfection of VSMCs with their wild-type expression vectors. Overexpression of Akt and STAT3 attenuated the inhibitory effect of empagliflozin on cyclin D (−41.3% versus +13.1% for Akt; −45.7% versus +4.4% for STAT3 overexpression, respectively, p<0.05), PCNA expression (−44.2% versus +12% for Akt; −37.6% versus +23.5% for STAT3 overexpression, respectively, p<0.05), and VSMC proliferation (−43.9% versus −17.5% for Akt; −52.7% versus +4.3% for STAT3 overexpression, respectively, p<0.05) (**Figures 3B-E**).

**Figure 3.**
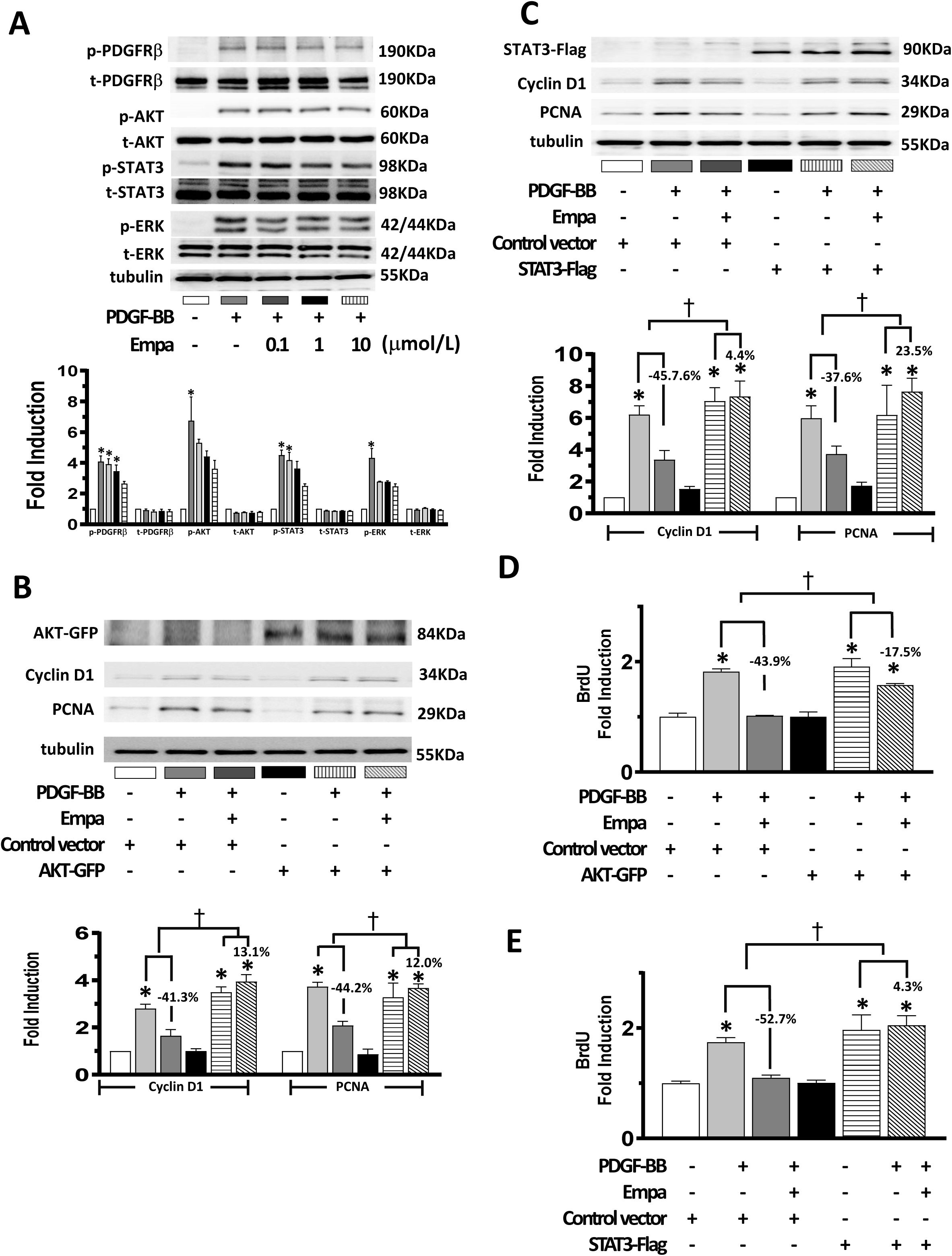
PDGF-related signaling in empagliflozin-inhibited VSMC proliferation. **A.** After 48 h of serum deprivation, VSMCs were treated with or without 60 ng/mL PDGF-BB and/or indicated concentrations of empagliflozin for 30 m. The expression of indicated proteins was evaluated by western blot (upper panels). The relative expression level of each protein was quantified by densitometry (lower panels). Each value represents mean ± SE of 4 independent experiments. Using one-way ANOVA, the different symbols (blank, *) represent the significant differences among groups. **B and C.** Growth-arrested VSMCs were pre-transfected with indicated plasmids for 24 h and subsequently treated with or without 60 ng/mL PDGF-BB and/or 10 μmol/L empagliflozin for 24 h. The expression of indicated proteins was evaluated by western blot (upper panels). The relative expression level of each protein was quantified by densitometry and normalized to the control level (lower panels). **D and E.** Growth-arrested VSMCs were pre-transfected with indicated plasmids for 24 h and subsequently treated with or without 60 ng/mL PDGF-BB and/or 10 μmol/L empagliflozin for 24 h. The proliferative activity of VSMCs was assayed by BrdU incorporation into VSMCs. Each value represents mean ± SE of 4 independent experiments. Using one-way ANOVA, the different symbols (blank, *) represent the significant differences among groups. Using two-way ANOVA, the symbol (†) represents the significant differences among groups.

We also evaluated whether empagliflozin could be involved in the migratory activity of VSMC, another crucial factor in determining the development of neointima formation.^5–7^ Consistently, empagliflozin treatment reduced PDGF-BB-induced VSMC migration. Furthermore, overexpression of Akt and STAT3 mitigated the inhibitory effect of empagliflozin on VSMC migration (−61.5% versus −38.5% for Akt; −61.5% versus −45% for STAT3 overexpression, respectively, p<0.05) (**Figure 4**). Collectively, these results imply that empagliflozin inhibits VSMC proliferation and migration via suppressing PDGF-BB-related signaling.

**Figure 4.**
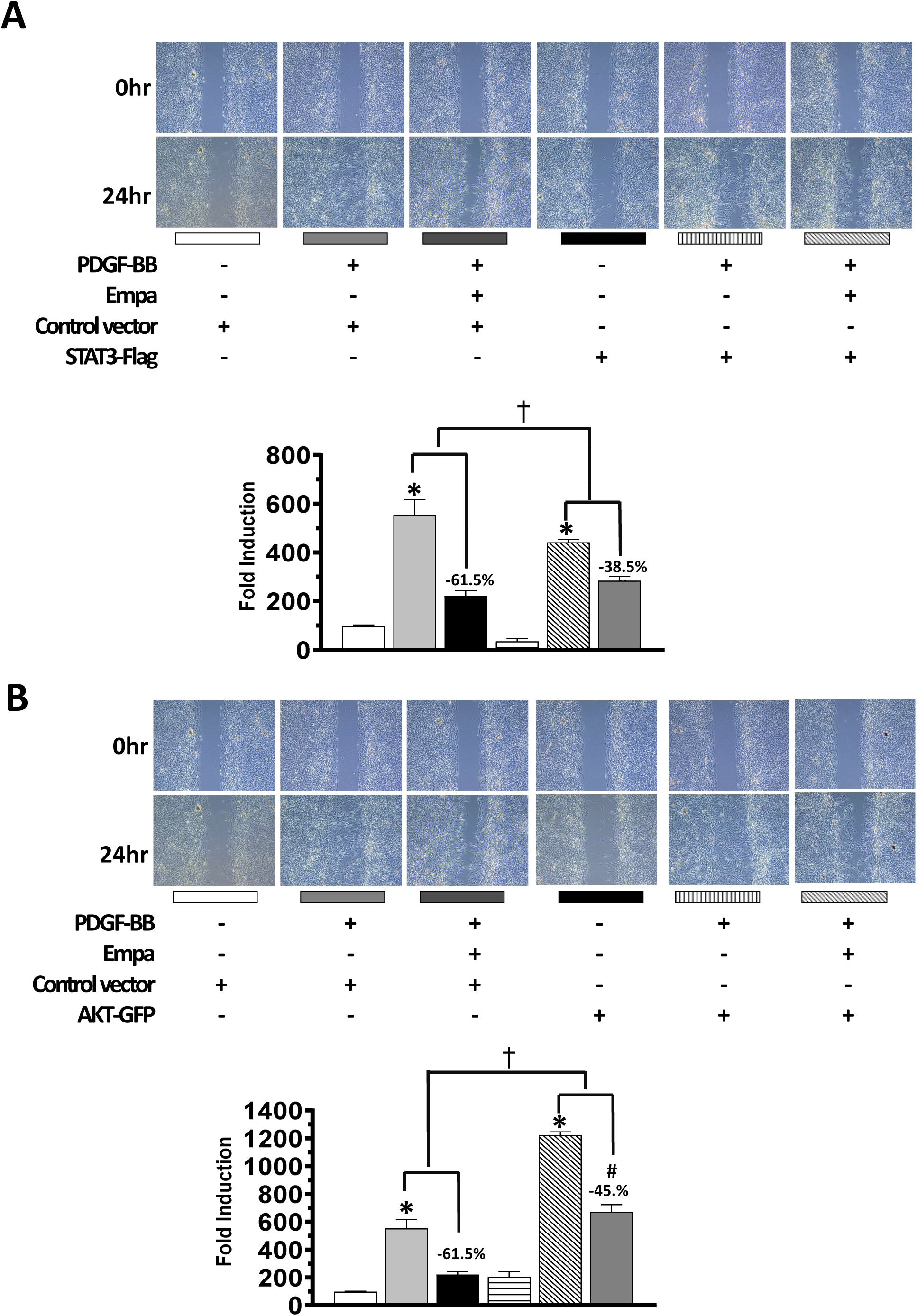
PDGF-related signaling in empagliflozin-inhibited VSMC migration. **A and B.** Growth-arrested VSMCs were pre-transfected with indicated plasmids for 24 h subsequently treated with or without 60 ng/mL PDGF-BB and/or 10 μmol/L empagliflozin for 24 h. The migratory activity of VSMC was assessed by wound healing assay. The protruded VSMCs were counted (magnification 100X) (upper panels). Each value (mean ± SE [n=6]) is determined by protruded cell number and normalized to the control level (lower panels). Using one-way ANOVA, the different symbols (blank, *, #) represent the significant differences among groups. Using two-way ANOVA, the symbol (†) represents the significant differences among groups.

### Direct effect of empagliflozin on VSMCs

Firstly, using RT/PCR, we observed that the expression of SGLT2 was minimal in either basal or PDGF-BB-treated rat VSMCs (**Supplemental Figure 2**), supporting prior reports that SGLT2 is only expressed in renal tubular cells.^15–17^ Furthermore, treatment of VSMCs with phlorizin, a non-specific SGLT inhibitor, did not influence the stimulating effects of PDGF-BB on VSMC proliferation and PDGF-related signaling expression (**Figure 5A**). Third, the inhibitory effects of empagliflozin on VSMC proliferation remained unchanged in SGLT2-deprived VSMCs (**Figures 5B and C**). Taken together, these findings offer further evidence that the effect of empagliflozin on VSMC functions is independent of SGLT2.

**Figure 5.**
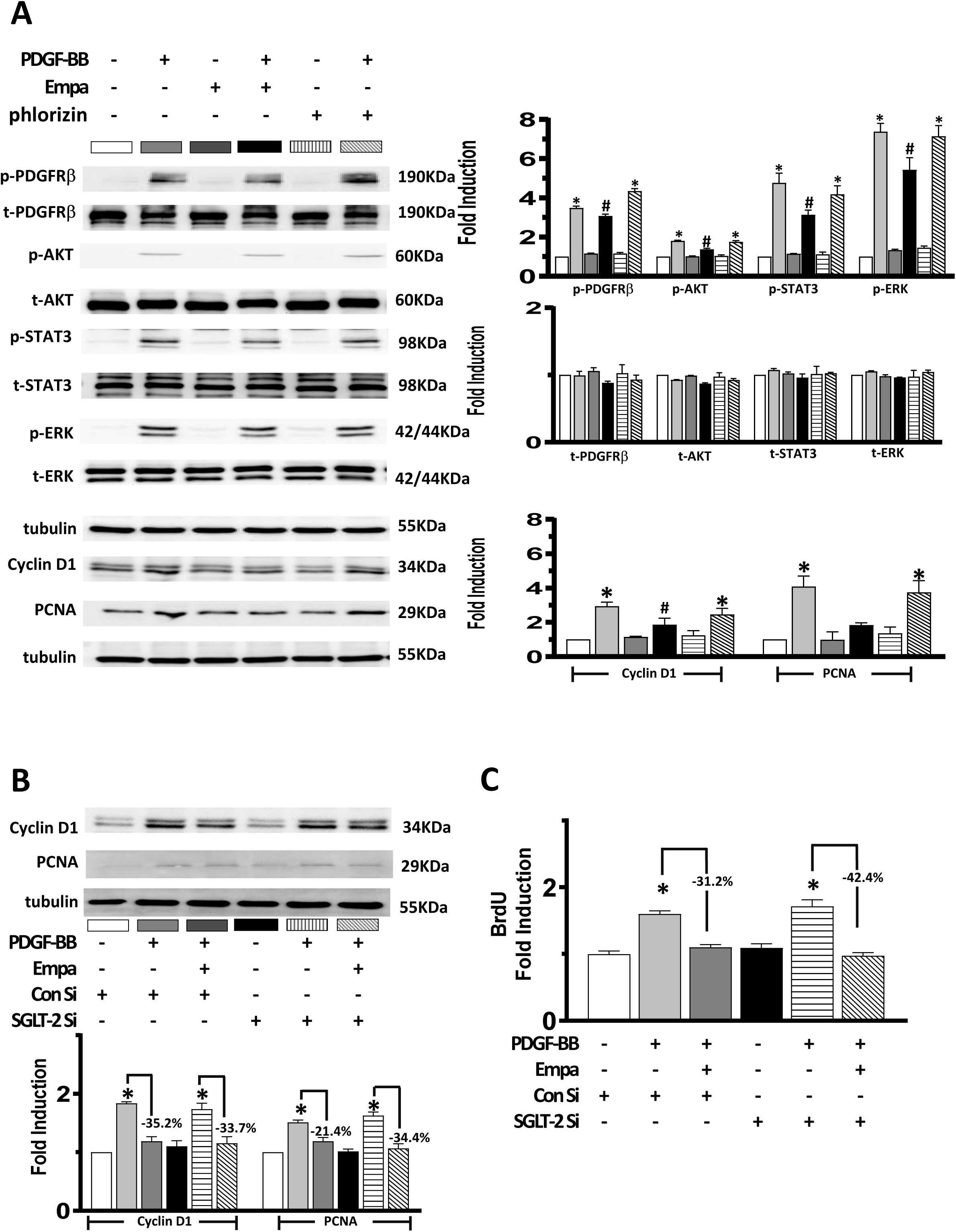
Direct effect of empagliflozin on VSMCs. **A.** After 48 h of serum deprivation, VSMCs were treated with or without 60 ng/mL PDGF-BB and/or indicated agents for 30 m to 24 h. The expression of indicated proteins was evaluated by western blot (left panels). The relative expression level of each protein was quantified by densitometry (right panels). Each value represents mean ± SE of 4 independent experiments. **B.** Growth-arrested VSMCs were pre-transfected with indicated siRNAs for 48 h and subsequently treated with or without 60 ng/mL PDGF-BB and/or 10 μmol/L empagliflozin for 24 h. The expression of indicated proteins was evaluated by western blot (upper panels). The relative expression level of each protein was quantified by densitometry and normalized to the control level (lower panels). **C.** Growth-arrested VSMCs were pre-transfected with indicated siRNAs for 48 h and subsequently treated with or without 60 ng/mL PDGF-BB and/or 10 μmol/L empagliflozin for 24 h. The proliferative activity of VSMCs was assayed by BrdU incorporation into VSMCs. Using one-way ANOVA, the different symbols (blank, *, #) represent the significant differences among groups. Using two-way ANOVA, the symbol (†) represents the significant differences among groups.

### Effect of empagliflozin on PDGF-related signaling expression *in vivo*

To confirm our *in vitro* data, we further determined the expression of PDGF-related signaling in the neointima induced by balloon injury. In consistent with the I/M area ratio changes, western blot indicated that the expression of p-Akt, p-STAT3, p-ERK, cyclin D, and PCNA was significantly higher in balloon-injured arteries than in control arteries (**Figure 6A**). In comparison with controls, their expressions were suppressed by empagliflozin treatment (**Figure 6A**). Immunohistochemical staining also partially documented the changes in western blot (**Figure 6B**).

**Figure 6.**
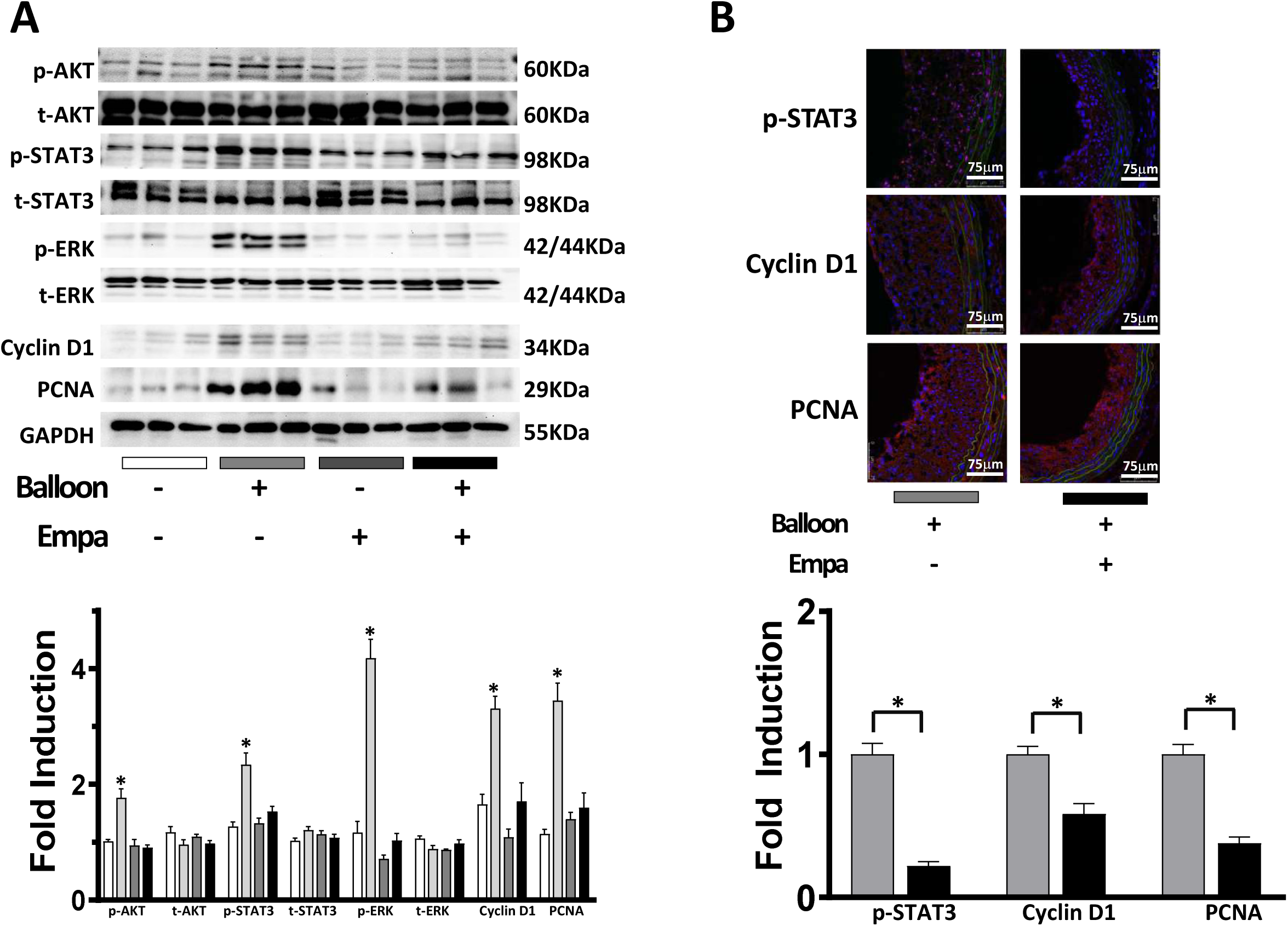
Effect of empagliflozin on PDGF-related signaling expression *in vivo*. **A.** The expression of indicated proteins was detected by western blot (upper panels). The relative expression level of each protein was quantified by densitometry and normalized to the control level (lower panels). Each value represents mean ± SE of 5 rats. Using one-way ANOVA, the different symbols (blank, *) represent the significant differences among groups. **B.** Representative confocal images show the expression of indicated proteins in the neointima of balloon-injured carotid arteries (upper panels). Labeling index of indicated proteins in the neointima was calculated and normalized to the control level (lower panels). Using unpaired t-test, the symbol (*) represents the significant difference between 2 groups.

## Discussion

The aim of this study is to evaluate whether empagliflozin (an SGLT2 inhibitor) exerts beneficial effects on VSMC functions. First, we observed that neointima formation was decreased in balloon-injured carotid arteries of empagliflozin-treated rats. Furthermore, empagliflozin might suppress PDGF-induced VSMC proliferation and migration, which was unrelated with cell apoptosis and cytotoxicity. Third, the inhibitory effects of empagliflozin on VSMC proliferation and migration were mediated via inactivation of PDGF-related signaling, such as PDGF-Rb, Akt, and STAT3. Finally, the inhibitory effect of empagliflozin on VSMC proliferation and migration is independent of its effects on SGLT2 and glucose metabolism.

Prior animal studies have demonstrated the protective effect of empagliflozin on neointima formation/atherosclerosis. Nevertheless, most such studies were performed in diabetic and/or hypercholesterolemic animals.^18–24^ To our knowledge, the present study is the first attempt to utilize non-diabetic rats for evaluating the effect of empagliflozin. Since empagliflozin treatment may not affect blood sugar levels in euglycemic animals,^15^ the inhibitory effect of empagliflozin on neointimal formation is obviously unrelated with its hypoglycemic action. Furthermore, we used an *in vitro* model to assess the direct effect of empagliflozin on cultured VSMCs. We disclosed that treating VSMCs with empagliflozin may suppress VSMC proliferation and migration, two key factors contributing to the development of neointimal formation.^5–7^ The *in vitro* results were verified by our *in vivo* rat model of balloon injury, which exhibited decreases of proliferative markers in the neointima after empagliflozin treatment. Taken together, our findings suggest the extra-glycemic effect of empagliflozin on inhibiting neointima formation.

SGLT2 is mainly expressed in the proximal renal tubule.^15–17^ Although some reports showed the minimal expression of SGLT2 in human VSMCs,^24, 25^ we demonstrated that SGLT2 is absent in rat VSMCs. In contrast, SGLT1 appears to be highly prevalent in cardiovascular tissues.^15, 16^ Since empagliflozin is highly selective for SGLT2 as compared with other SGLT inhibitors,^15^ the contributing role of SGLT2 in the inhibitory effect of empagliflozin on neointimal formation could be virtually excluded. Furthermore, treatment of VSMCs with phlorizin, a non-specific SGLT inhibitor,^15^ did not display a similar response as empagliflozin. Finally, silencing SGLT2 did not alter the effects of empagliflozin of VSMC functions. These findings further supported the SGLT2-independent effect of empagliflozin. Our findings conflict with those of a prior report showing the involvement of SGLT2 in empagliflozin-induced inhibition of VSMC proliferation.^25^ The differences in spices, agonists used, and study design may explain this disparity.

The involvement of PDGF and PDGFR (especially PDGF-BB and PDGF-Rb) in the pathogenesis of neointimal formation has been a subject of great interest. This notion is based on prior reports showing the greater activation of PDGF and PDGFR-related signaling in the neointima.^9, 10, 26^ Furthermore, blocking PDGF/PDGFR-related signaling, such as PDGF antibodies, PDGF-BB aptamers, kinase inhibitors, and PDGF-Rb neutralizing antibodies, has been demonstrated to reduce neointimal formation in animal models.^9, 10, 26^ The novel finding in this study is to disclose the suppressive effects of empagliflozin on PDGFR-related signaling, including phosphorylation of PDGF-Rb, Akt, and STAT3. Furthermore, overexpression of PDGFR-related signaling attenuated the inhibitory effect of empagliflozin on VSMC proliferation. These results suggest that suppressing PDGF-related signaling may contribute to the inhibitory effect of empagliflozin on VSMC proliferation. Beyond inhibiting SGLT2, empagliflozin has been also emerged as an inhibitor of PDGFR-related signaling. Because empagliflozin did not induce cell apoptosis and cytotoxicity, it may be an appropriate agent for preventing or treating vascular diseases involving VSMC proliferation in clinic applications.

SGLT2 inhibitors may possess other effects that contribute to their protective effects on the development of neointimal formation/atherogenesis, including suppressing vascular inflammation and oxidative stress, improving endothelial function, reducing foam cell formation, and preventing platelet activation.^16, 17, 27^ However, how these other effects contribute to the reductions in VSMC proliferation and neointimal formation during empagliflozin treatment merits further investigation. Nevertheless, the present study provided novel findings about the direct effects of empagliflozin on VSMC functions.

The maximal concentration (10 μmol/L) of empagliflozin we applied *in vitro* exceeded the blood concentration measured in humans given a standard dose with 25 mg/kg/day (Cmax between 0.4-1.1 μmol/L).^16, 17, 28^ Nevertheless, treating VSMCs with a therapeutic concentration (1 μmol/L) of empagliflozin may still have some effects, even if we observed no such significant difference. An application of higher concentration *in vitro* is a common approach for evaluating acute effects and predicting the long-term effects of these agents on human disorders. Furthermore, the blood concentration of empagliflozin measured in balloon-injured rats was within the therapeutic range. Conceivably, empagliflozin may be employed to suppress VSMC proliferation and neointimal formation in clinical practice.

This study has some limitations. First, whether the findings regarding empagliflozin could be applied to other SGLT2 inhibitors warrants further study. Furthermore, the application of our findings from rats to humans is still in question. Third, the effect of empagliflozin on other functions of VSMCs, such as extracellular matrix production and differentiation, that also contribute to the development of neointimal formation requires further investigation. Fourth, further study is needed to determine the exact working site at which empagliflozin participates in PDGF-related signaling. Finally, although empagliflozin is highly selective for SGLT2, the contributing role of SGLT1 in empagliflozin-related effects still could not be excluded.

In conclusion, this study highlights the crucial role of PDGF-related signaling in mediating the inhibitory effects of empagliflozin on VSMC proliferation and/or neointimal formation, which are unrelated with its effects on SGLT2 and glucose metabolism. These findings provide an additional mechanism explaining for the clinical benefit of empagliflozin in preventing cardiovascular events.

## Acknowledgement

None

